# Glucose Metabolism Sustains Aberrant STAT3 Signaling in Colorectal Cancer via Glycosylated Paracrine Factors

**DOI:** 10.1101/2025.07.22.666143

**Authors:** Kathryn Buscher, Kelsey Temprine, Christopher Mays, Noora Aabed, Samuel Kerk, Hannah N. Bell, Joseph A. Nieto Carrion, Harrison Greenbaum, Varun Ponnusamy, Sadeesh K. Ramakrishnan, Costas A. Lyssiotis, Xiang Xue, Yatrik M. Shah

## Abstract

The JAK-STAT3 signaling pathway is a key driver of colorectal cancer (CRC) progression. While STAT3 is canonically activated by cytokines such as IL-6, this activation is typically transient due to negative feedback mechanisms. In CRC, however, STAT3 is aberrantly and persistently activated, promoting tumor cell proliferation and survival. Here, we demonstrate that glucose sustains STAT3 activation independent of cytokine availability. By manipulating glucose metabolism, we show that both glucose and its downstream metabolite, GlcNAc, are essential for maintaining STAT3 activation. Moreover, cells with high basal STAT3 activity produce glucose-dependent glycosylated proteins that can activate STAT3 in neighboring cells via paracrine signaling. Proteomic analysis identified multiple candidate proteins involved in this process; however, no single protein was sufficient to fully activate STAT3, suggesting that a combination of glycosylated proteins likely acts synergistically. In vivo, inhibition of glycolysis reduces STAT3 activation in tumors, and genetic deletion of STAT3 in subcutaneous tumor models significantly decreases tumor growth. Together, these findings uncover a novel mechanism by which glucose metabolism supports sustained STAT3 activation in CRC, highlighting a potential metabolic vulnerability for therapeutic targeting.

## INTRODUCTION

Colorectal cancer (CRC) is the third most common cancer in the United States, ranking third and fourth in mortality for males and females, respectively^1^. Although CRC incidence has declined over the past two decades due to screening programs, early-onset CRC, defined as diagnosis before 49 years of age, had been rising and now constitutes nearly 20% of all new CRC diagnosis^2^. The genomic landscape of CRC is characterized by a well-defined sequence of mutations that drive tumor initiation and progression. APC mutations occur early and are found in approximately 80% of sporadic CRC cases^3^. APC encodes a critical negative regulator of the Wnt signaling pathway; its loss leads to constitutive Wnt activation, resulting in uncontrolled epithelial proliferation and adenoma formation. Subsequent activating mutations in KRAS, observed in approximately 40% of cases, promote adenoma growth by constitutively activating the RAS-RAF-MEK-ERK signaling cascade, enhancing cell proliferation and survival. As the disease progresses, inactivation of the tumor suppressor TP53, present in more than 50% of CRCs facilitates the transition from benign adenoma to invasive carcinoma^4^. These core mutations are frequently retained in metastatic lesions, which remain largely refractory to current therapies and are a major contributor to CRC-associated mortality^3^. Despite advances in surgical resection and adjuvant therapies, metastatic CRC continues to have poor prognosis, highlighting the urgent need for novel therapeutic strategies^5^.

The Janus kinase (JAK)/signal transducer and activator of transcription 3 (STAT3) signaling pathway is emerging as a promising therapeutic target in CRC^6^. Canonical STAT3 activation is initiated by a cytokine, such as interleukin-6 (IL-6), binding to its receptor leading to JAK protein recruitment. JAK proteins recruit STAT3, which is then phosphorylated, leading to dimerization, and translocation to the nucleus to regulate transcription of genes promoting proliferation and survival^6,7^. Genetic disruption of IL-6 or enterocyte-specific STAT3 markedly reduces tumor burden in murine models, underscoring the oncogenic role of the IL-6-STAT3 signaling axis^8^.

Under physiological conditions, cytokine-induced JAK-STAT3 signaling is transient and subject to negative feedback regulation by suppressor of cytokine signaling 3 (SOCS3)^9,10^. However, sustained STAT3 activation has been shown to be critical for tumor proliferation across several CRC models^8,11–13^. The mechanisms underlying persistent STAT3 activation remain poorly understood.

Targeting the JAK-STAT3 pathway has been explored as a potential therapeutic approach for CRC. Although several JAK and STAT3 inhibitors have undergone clinical trials, none are currently approved for CRC treatment^6,14^. Notably, JAK inhibitors are used to treat other cancers, including lung and breast cancers^15^. Elucidating the mechanisms sustaining STAT3 activation could pave the way for the development of more effective inhibitors in CRC.

Here, we identify glucose metabolism as a key regulator of STAT3 activation in CRC, independent of cytokine stimulation. Nutrient deprivation experiments revealed that glucose is essential to sustain both basal and cytokine-induced STAT3 activation. Further analyses demonstrated that STAT3 activation depends on the production of glycosylated secreted proteins that promote autocrine and paracrine STAT3 signaling. Our data suggests that multiple secreted factors act synergistically to sustain STAT3 signaling. Proteomic analyses supported this hypothesis by revealing several candidate secreted proteins. In vivo, genetic deletion of STAT3 markedly reduced tumor size, and pharmacological inhibition of glycolysis diminished STAT3 activation in tumors.

Collectively, these findings delineate a previously unrecognized role for glucose metabolism in sustaining oncogenic STAT3 signaling in CRC and suggest that targeting glucose metabolism for STAT3 inhibition represents a novel therapeutic strategy.

## RESULTS

### STAT3 activation in CRC is glucose dependent

CRCs often exhibit high levels of basal STAT3 activation, a key driver of tumor progression and survival^8,11–13^. Given the importance of STAT3 in CRC, understanding how its activation is sustained despite the presence of strong negative feedback regulators is critical.

Characterization of a panel of human CRC cell lines revealed that several exhibited high levels of basal STAT3 activation (Fig. 1A). We hypothesized that this could represent an important mechanism to sustain STAT3 activity in CRC, even in the presence of strong negative feedback regulators. To investigate the mechanism driving this activation, we began by systematically evaluating the role of nutrients in the culture media. In cells cultured in DMEM supplemented with 10% FBS, STAT3 activation was unaffected by serum deprivation (Supp. Fig. 1A).

**Figure 1.**
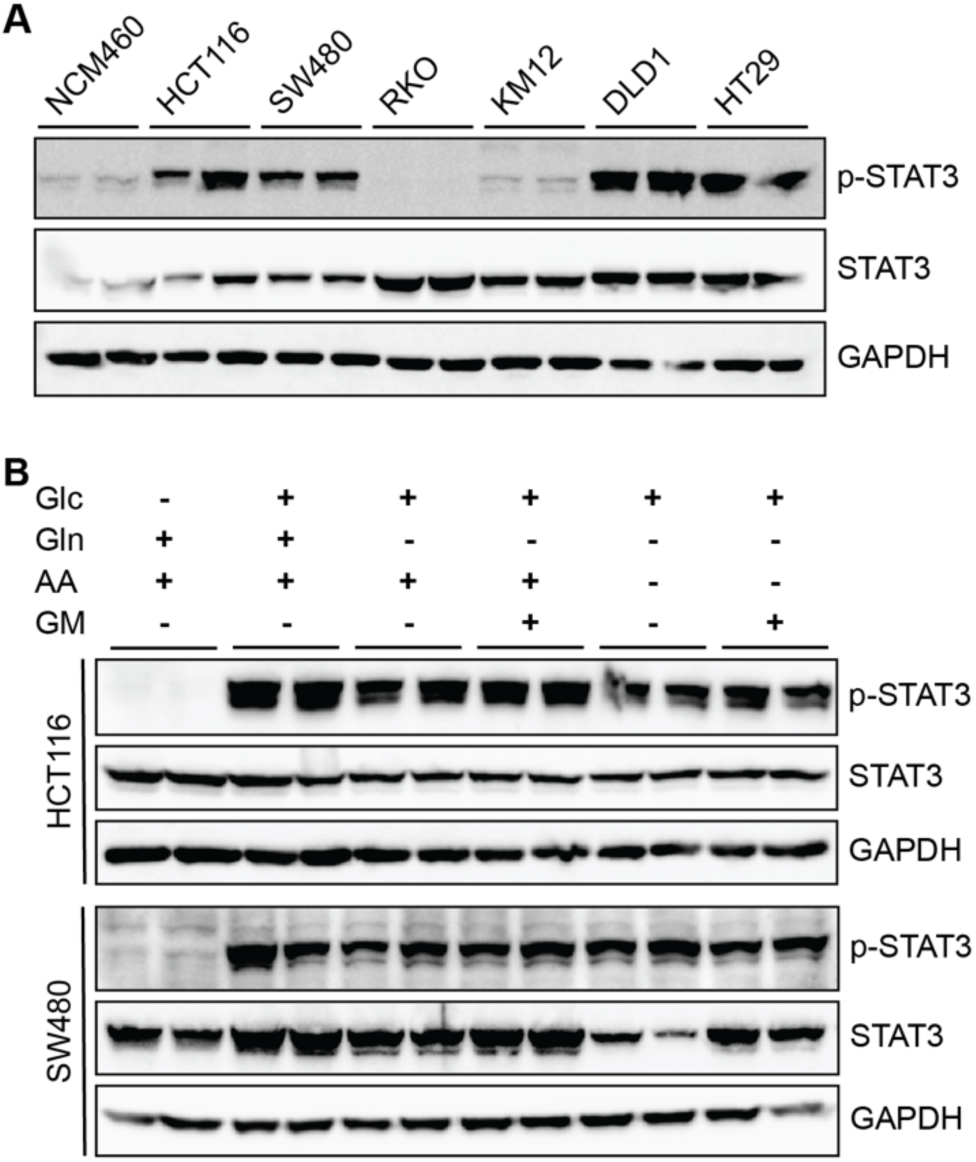
STAT3 activation is regulated by glucose in colorectal cancer. (A) Normal colon epithelium (NCM460) or colorectal cancer (CRC) derived (HCT116, SW480, RKO, KM12, DLD1, HT29) cell line lysates probed for basal phosphorylated STAT3 and total STAT3 levels with GAPDH loading control. (B) CRC cell lines were subjected to glucose (Glc), glutamine (Gln), amino acid (AA), or glutamax (GM) drop out for 24 hours prior to harvesting lysates and probing for phospho-STAT3 and total STAT3 levels with GAPDH loading control.

Similarly, removing glutamine or essential amino acids did not alter STAT3 activation. However, glucose deprivation selectively and significantly reduced STAT3 activation (Fig. 1B).

Consistent with these findings, glucose dependence of STAT3 activation was also observed in pancreatic, liver, and cervical cancer cell lines (Supp. Fig. 1B). Interestingly, glucose specifically induced STAT3 activation, whereas other signaling pathways, such as AKT and MAP kinase, were either unaffected or downregulated under the same conditions (Supp. Fig. 1C). Gene expression analysis of canonical STAT3 signaling components revealed no significant differences between glucose-replete and glucose-depleted conditions (Supp. Fig. 1D). These results suggest that glucose-mediated STAT3 activation occurs independently of transcriptional regulation of canonical upstream components or negative feedback regulators, such as SOCS3^9,10^.

### Physiological glucose levels sustain basal and cytokine-induced STAT3 activation

Cancer cells are characterized by high glucose consumption and are typically exposed to glucose concentrations of 4-6mM in circulating blood. In contrast, standard cell culture media often contains 25mM glucose. To assess whether glucose concentrations within the physiological range can activate STAT3, CRC cells were exposed to glucose levels ranging from 0 to 25 mM. STAT3 activation peaked at 5 mM glucose, which closely mirrors physiological blood glucose levels^16^ (Fig. 2A). Time-course experiments demonstrated that STAT3 activation began within 1-2 hours of glucose exposure, reaching its maximum at 24 hours (Fig. 2B). This gradual activation suggests a non-canonical mechanism, distinct from the rapid cytokine-induced STAT3 signaling observed in other contexts^17,18^.

**Figure 2.**
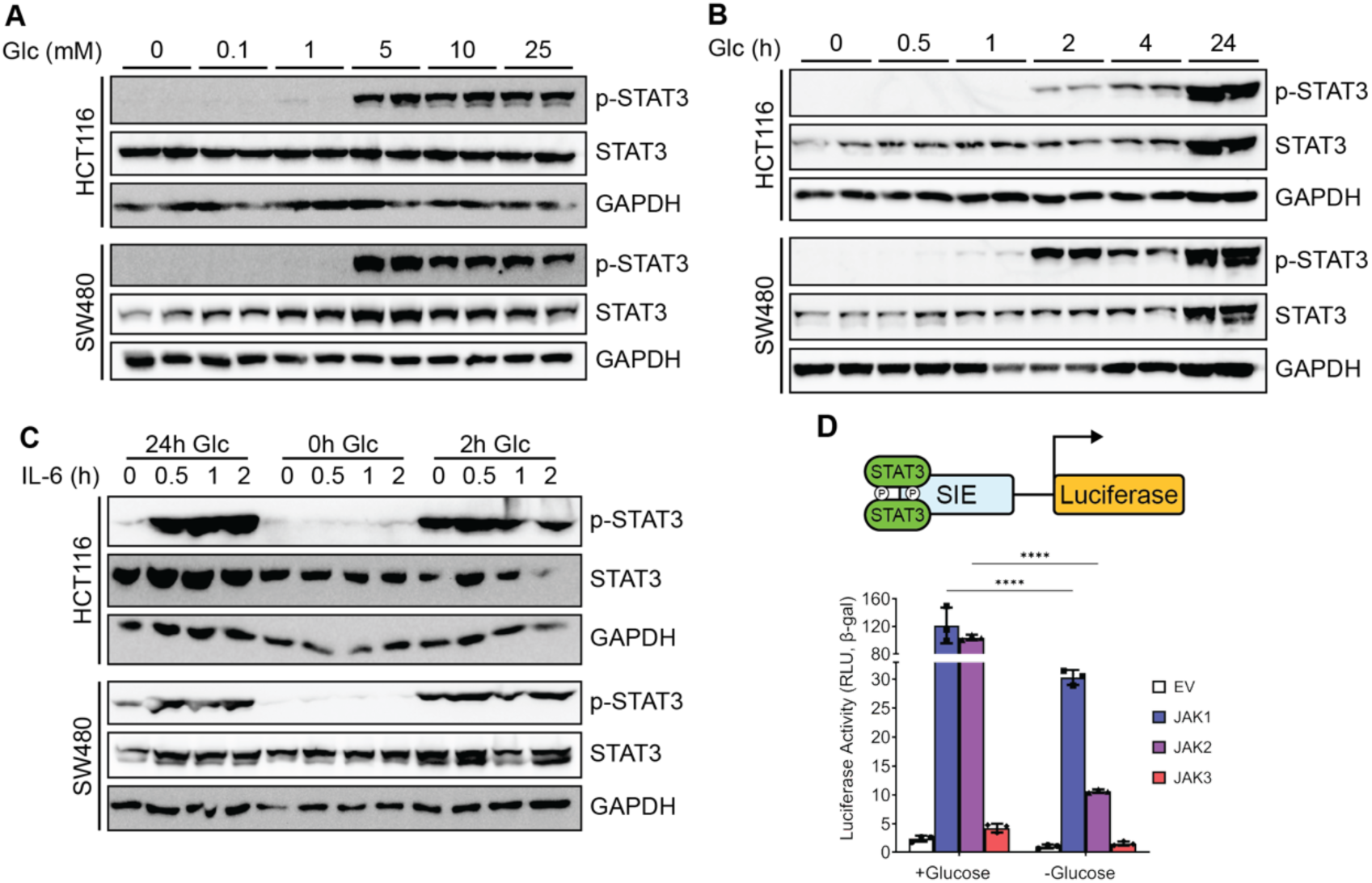
Glucose is required to maintain basal STAT3 activation and synergizes with canonical IL-6-JAK signaling pathways. (A) HCT116 and SW480 cell lines were treated with increasing concentrations of glucose for 24 hours prior to harvesting lysates and probing for phospho-STAT3 and total STAT3 levels with GAPDH loading control. (B) HCT116 and SW480 cell lines were glucose deprived overnight then treated with 25mM glucose and assessed for phospho-STAT3, total STAT3, and GAPDH in a temporal manner. (C) HCT116 and SW480 cell lines treated with glucose (25mM) for 0, 2, or 24 hours and IL-6 (10ng/mL) for 0, 0.5, 1, or 2 hours prior to harvesting lysates and probing for phospho-STAT3 and total STAT3 levels with GAPDH loading control. (D) Top: Schematic of the luciferase assay performed. Bottom: Results of luciferase assay done in HEK293T cells co-transfected with JAK overexpression constructs. Cells were treated with 0mM or 25mM glucose for 24 hours.

In CRC, IL-6 is a major mechanism of STAT3 activation^8^. To determine whether glucose is essential for IL-6-mediated STAT3 activation, cells were treated with IL-6 in the presence or absence of glucose. Under glucose-deprived conditions, IL-6 failed to activate STAT3 (Fig. 2C). Canonical JAK-STAT3 signaling involves cytokine receptor activation, leading to JAK phosphorylation, which in turn phosphorylates and activates STAT3^6,7^. To investigate the role of JAK proteins in glucose-mediated STAT3 activation, cells were treated with the JAK inhibitor, Ruxolitinib^19^. Inhibition of JAK signaling with Ruxolitinib completely abolished glucose-mediated STAT3 activation (Supp. Fig. 2). To further explore the involvement of JAK proteins, we used a sis-Inducible Element (SIE)-luciferase reporter, which luminesces following STAT3 activation.

Overexpression of JAK1 or JAK2 did not fully restore STAT3 activation in glucose-depleted cells (Fig. 2D). Collectively, these findings indicate that canonical STAT3 activation pathways require glucose and that JAK proteins are integral to glucose-mediated STAT3 activation.

### A glycolytic intermediate plays a role in glucose mediated STAT3 signaling

To define the role of glucose metabolism in STAT3 activation, we utilized the glycolytic inhibitor 2-deoxy-glucose (2-DG), which blocks glycolysis by targeting hexokinase (Fig. 3A). Treatment with 2-DG markedly reduced STAT3 activation, even in the presence of IL-6 (Fig 3B-C), indicating that glycolytic activity is essential for this process. To identify the step in the glycolytic pathway required for STAT3 activation, we supplemented cells with pyruvate, the product of glycolysis. However, neither pyruvate, nor its cell-permeable analog, methyl-pyruvate restored STAT3 activation in glucose-deprived conditions, but p-ACC and p-AMPK both increased with pyruvate addition (Fig. 3D-E, Supp. Fig. 3A). Since pyruvate is rapidly converted to lactate in tumor cells, we next investigated whether lactate metabolism is essential for STAT3 activation. Inhibition of lactate dehydrogenase (LDH) using Galloflavin did not attenuate STAT3 activation (Supp. Fig. 3B). Additionally, hypoxic conditions decrease STAT3 activation in the absence of glucose in a short time frame, but do not affect STAT3 activation when glucose is not depleted (Supp. Fig. 3C). These findings collectively suggest that STAT3 activation relies on a specific glycolytic intermediate, upstream of pyruvate, highlighting a critical and previously unappreciated metabolic dependency for this signaling pathway.

**Figure 3.**
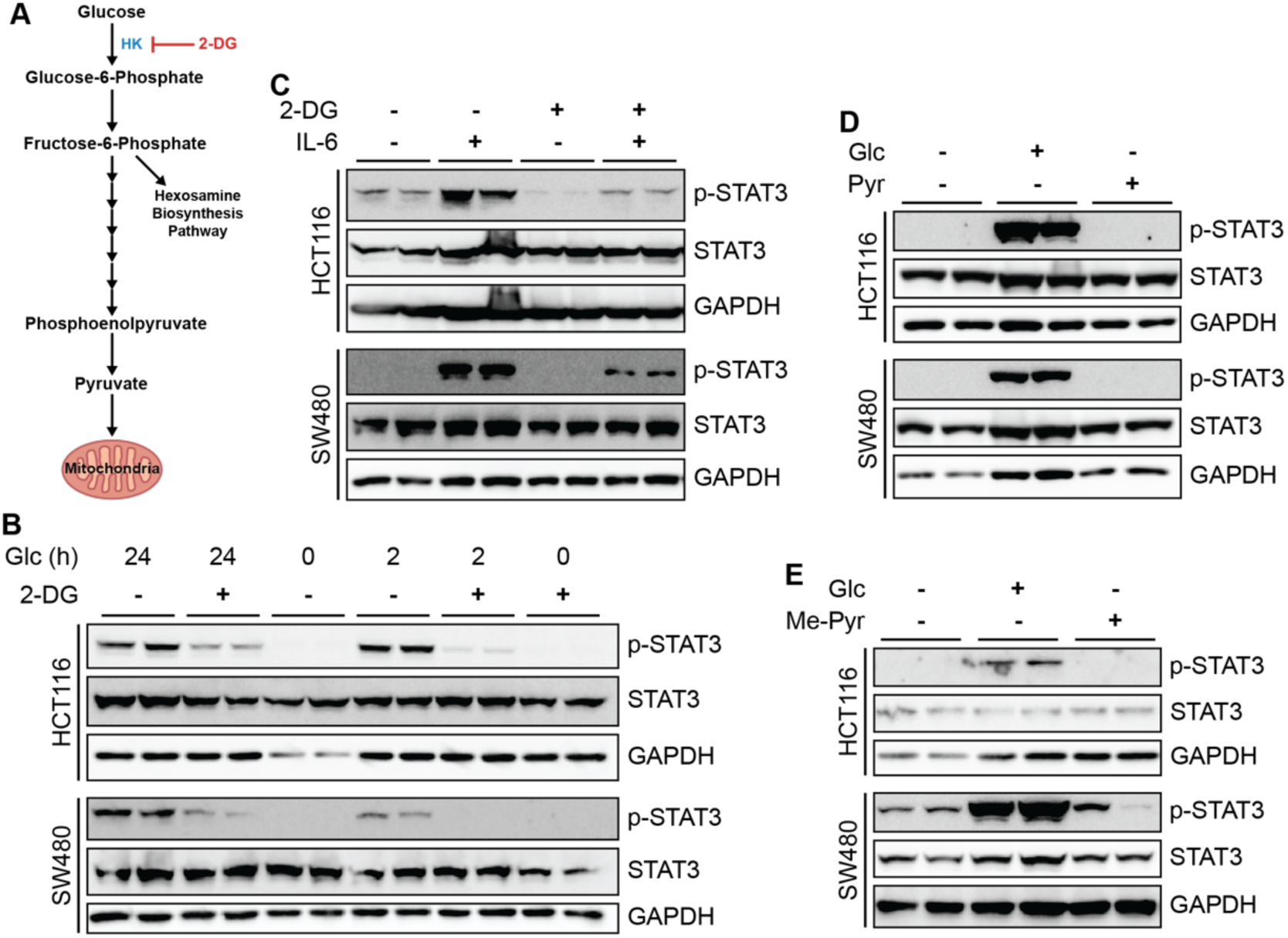
A glycolytic intermediate plays a role in maintaining STAT3 activation. (A) Diagram of glycolysis showing inhibition of hexokinase (HK) with 2-deoxyglucose (2-DG). (B) CRC cell lines treated with glucose (25mM) for 0, 2, or 24 hours and 2-DG (2.5mM) for 2 hours prior to harvesting lysates and probing for phospho-STAT3 and total STAT3 with GAPDH loading control. (C) CRC cell lines treated with 2-DG (2.5mM) for 2 hours and IL-6 (10ng/mL) for 2 hours. (D) CRC cell lines treated with glucose (25mM) or sodium pyruvate (25mM) overnight. (E) CRC cell lines treated with methyl-pyruvate (50mM) or glucose (25mM) for 4 hours.

### Glycosylation is important for glucose mediated STAT3 activation

To comprehensively investigate glucose-mediated pathways, bulk RNA sequencing was performed. Cells were deprived of glucose overnight, followed by glucose reintroduction for 4 hours to identify early gene expression changes upon glucose supplementation. The primary pathway upregulated was associated with a gene signature indicative of endoplasmic reticulum (ER) stress^20,21^ (Fig. 4A, Supp. Table 1). It is previously well established that removing glucose from cells increases ER stress^22–24^. Consistent with this, glucose deprivation increased BIP expression, a marker of ER stress^20,21^ (Supp. Fig. 4A). To determine whether ER stress contributes to STAT3 activation, cells were treated with two canonical ER stress inducers.

**Figure 4.**
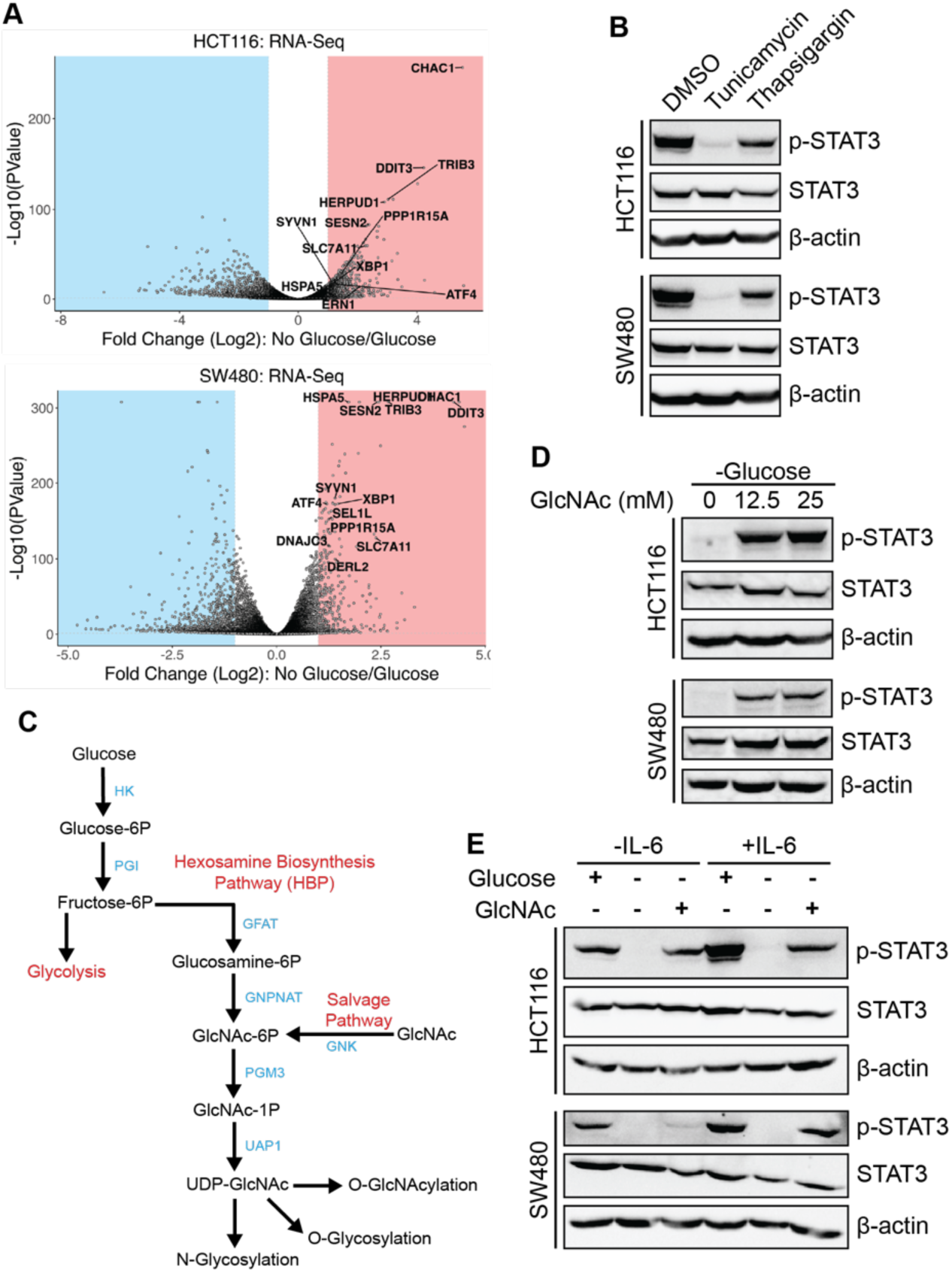
GlcNAc can activate STAT3 when glucose is depleted. (A) Volcano plots for RNA-Seq data from CRC cell lines treated with 0 or 25mM glucose for 4 hours. ER stress genes are labelled in the plots. (B) CRC cell lines treated with ER stress inducers tunicamycin (6µM) or thapsigargin (2µM) or DMSO control for 24 hours. (C) Schematic of the hexosamine biosynthesis pathway (HBP). (D) CRC cell lines treated with glucose free media overnight with 0, 12.5, or 25mM N-acetylglucosamine (GlcNAc) for 24 hours. (E) CRC cell lines treated with 0 or 25mM glucose for 24 hours, GlcNAc (25mM) for 24 hours, and IL-6 (20ng/mL) for 30 minutes.

**Table 1.**
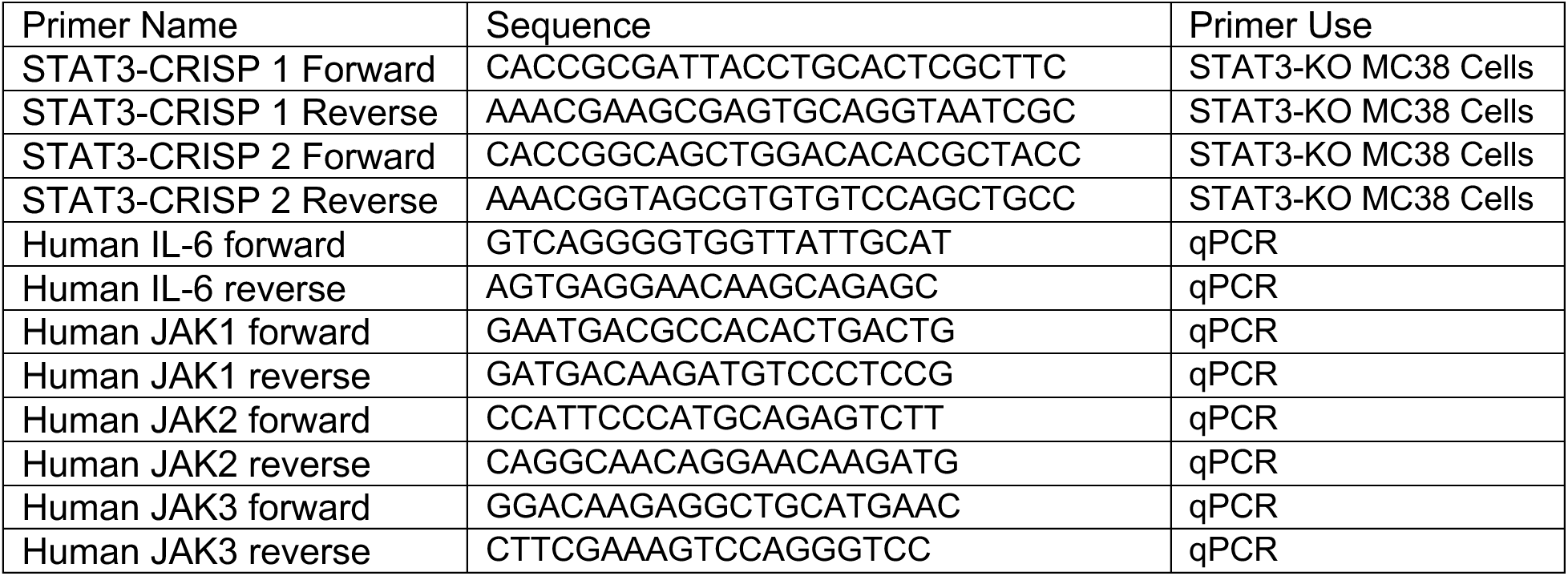
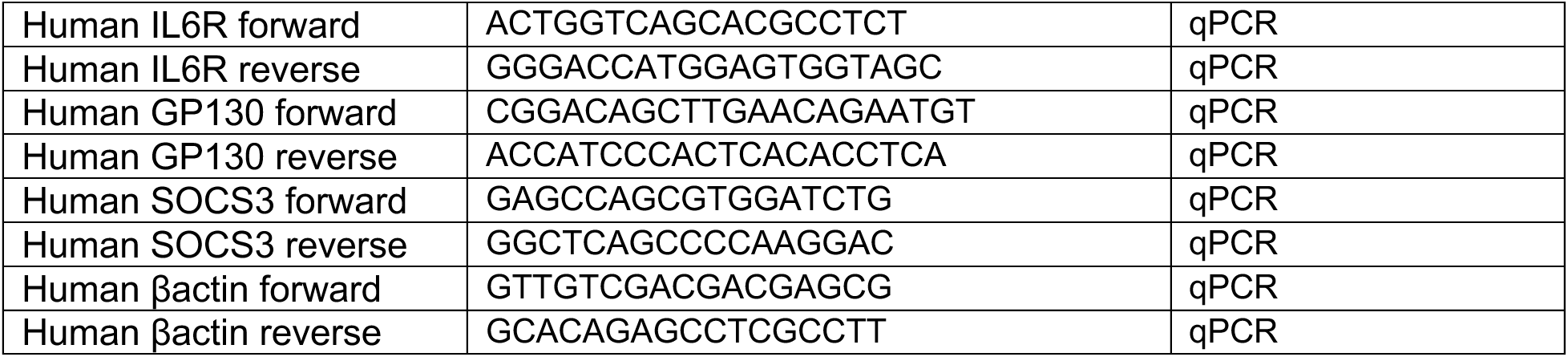
Primer Sequences.

STAT3 activation was reduced by thapsigargin, which induces ER stress through calcium homeostasis disruption^25^. However, tunicamycin, an inhibitor of N-glycosylation, nearly abolished STAT3 activation^26^ (Fig. 4B).

Glucose plays a critical role in protein glycosylation via the hexosamine biosynthesis pathway (HBP), which branches off glycolysis at fructose-6-phosphate to produce uridine diphosphate N-acetylglucosamine (UDP-GlcNAc) (Fig. 4C). To confirm the involvement of the HBP and glycosylation in STAT3 activation, cells were treated with GlcNAc, which directly enters the HBP and sustains glycosylation independently of glucose. GlcNAc supplementation restored STAT3 activation in glucose-depleted cells, both in the presence and absence of IL-6 (Fig. 4D-E). Further analysis revealed that inhibition of O-glycosylation with OSMI-1 had no effect on STAT3 activation^27^ (Supp. Fig. 4B). In contrast, inhibition of N-glycosylation with NGI-1 significantly reduced STAT3 activation^28^ (Supp. Fig. 4C). Other monosaccharides can substitute for glucose and enter the HBP^29^. Among monosaccharides tested, mannose, which is critical for N-glycosylation, showed the highest activation levels^30^ (Supp Fig. 4D-E). Collectively, these results highlight the importance of N-glycosylation in sustaining STAT3 activation, underscoring a critical link between glucose metabolism and STAT3 signaling.

### A glucose-dependent secreted protein activates STAT3

Glycosylation is a common post-translational modification essential for protein secretion^31^. To investigate if a secreted protein is required for STAT3 activation, cells were treated with Brefeldin A (BFA), an inhibitor of vesicle transport between the ER and Golgi, or GolgiStop (GS), which blocks secretion via the Golgi^32,33^. Both treatments led to a reduction in STAT3 activation (Fig. 5A). Moreover, conditioned media from cell lines with high basal STAT3 activation (HCT116 and SW480) increased STAT3 activation in RKO cells and NCM460 (normal epithelium) cells, which have low basal STAT3 activation (Fig. 5B and Supp Fig. 5A). This finding indicates that a secreted factor plays a key role in STAT3 activation through autocrine or paracrine signaling. Boiling conditioned media from HCT116 or SW480 cells reduced STAT3 activation (Fig. 5C), and CM filtered to <30 kDa failed to activate STAT3 (Supp. Fig. 5B), indicating that the factor is heat-labile and likely a protein.

**Figure 5.**
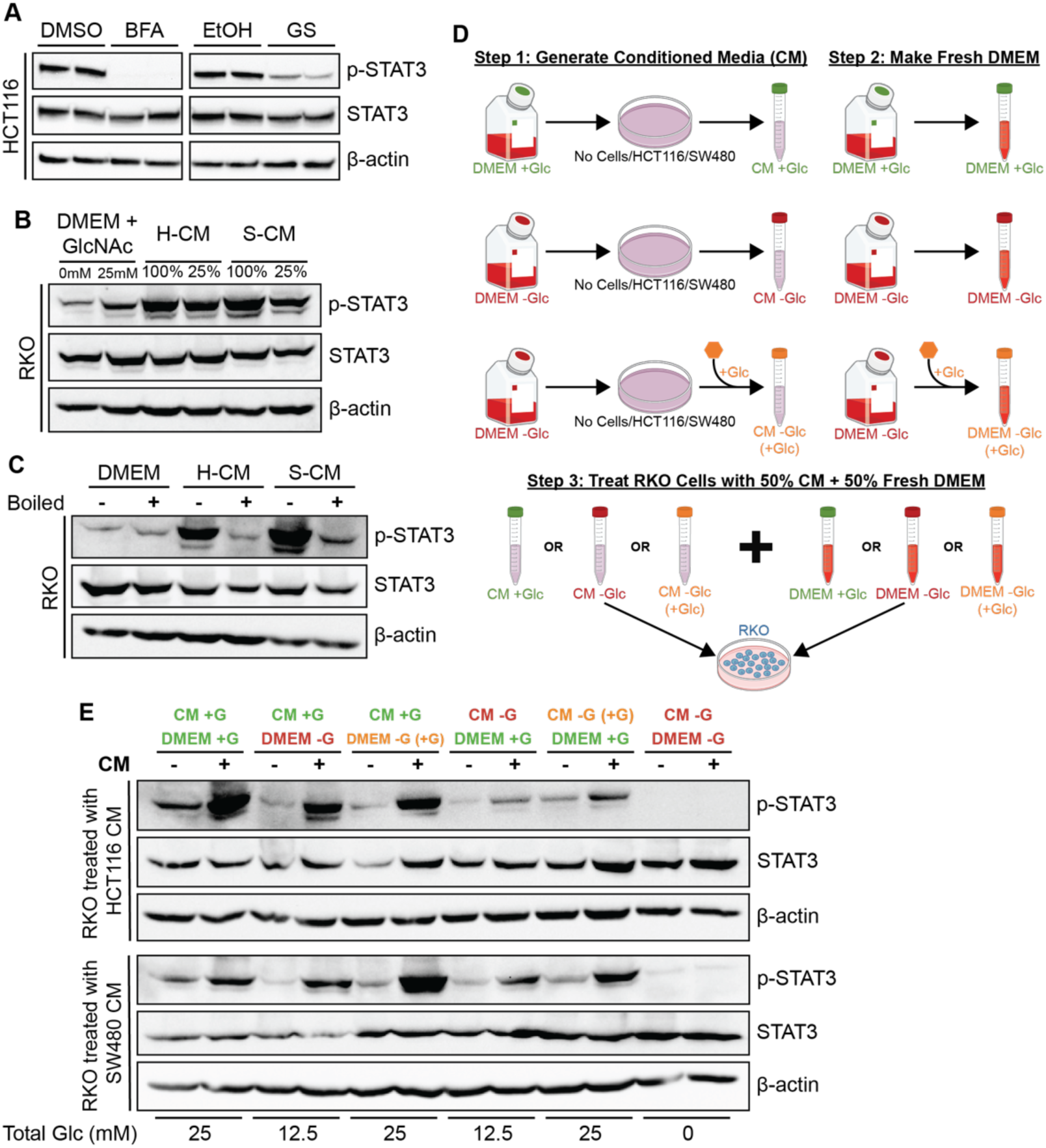
Glucose-mediated STAT3 activation occurs through a secreted protein. (A) CRC cell lines treated with brefeldin A (BFA; 10ng/mL) or golgi stop (GS; 1:1000 dilution) for 5 hours. (B) RKO cells treated with 0 or 25mM GlcNAc control, HCT116 condition media (H-CM), or SW480 condition media (S-CM) for 48 hours with refreshed CM at 24 hours. RKO cells were treated with either pure condition media or condition media diluted with DMEM (25% CM). (C) RKO cells treated with boiled (incubated at 100°C for 10 minutes) or non-boiled DMEM, HCT116 condition media (H-CM), or SW480 condition media (S-CM) for 48 hours with refreshed CM at 24 hours. (D) Schematic of condition media experiment performed in E. (E) RKO cells treated with HCT116 or SW480 CM or mixed DMEM used on HCT116, and SW480 cell lines final concentrations of glucose are 0, 12.5, or 25mM. RKO was treated with CM for 48 hours with refreshed CM at 24 hours.

To determine the role of glucose in the production of the secreted protein, conditioned media was collected from cells cultured in the presence and absence of glucose. In some conditions, glucose was reintroduced after conditioned media was generated to assess if glucose is required to produce the secreted factor, its ability to elicit a response, or both. (Fig. 5D). We found that glucose was essential to produce the secreted protein capable of activating STAT3. Conditioned media generated in the absence of glucose failed to fully activate STAT3, and reintroducing glucose after media collection did not fully rescue this defect (Fig. 5E). These results demonstrate that glucose is required for the production of the secreted protein that drives STAT3 activation, consistent with a model in which glucose-dependent glycosylation is critical for its function.

### Investigation of secreted protein candidates

To identify secreted protein candidates, quantitative mass spectrometry with tandem mass tagging was performed on conditioned media from HCT116, SW480, and RKO cells (Fig. 6A). The fold change in protein abundance between HCT116 or SW480 cells and RKO cells was calculated (Fig. 6B). A total of 77 proteins were upregulated in both HCT116 and SW480 cells. Additionally, we performed a meta-analysis focused on proteins that are secreted, glycosylated, altered in cancer, and function as bona fide activators of STAT3 defined by their ability to induce STAT3 activation when overexpressed and suppress activation when knocked down. From our 77 upregulated proteins, 23 have previously been implicated in activating STAT3 (Supp. Table 2). One protein if interest, CD109, is a glycosylphosphatidylinositol-anchored glycoprotein whose expression is elevated in multiple cancer types^34–36^. Increased CD109 expression is associated with tumor progression and poor prognosis^34,35^. Prior studies have implicated CD109 in the regulation of several signaling pathways, including EGFR and STAT3 signaling^34–39^. In various cancer cell lines, STAT3 activation is diminished following CD109 knockout, whereas CD109 overexpression enhances STAT3 signaling^37–39^. Additionally, Filppu et al. demonstrated that CD109 can bind glycoprotein 130 (GP130), a core component of the IL-6 receptor complex^37^. Despite these observations, recombinant CD109 has not been shown to directly activate STAT3 in vitro, and it remains unclear whether metabolic cues such as glucose availability influence CD109 expression. We found that endogenous CD109 expression is sensitive to glucose levels, with reduced glucose availability leading to decreased CD109 expression (Fig. 6C). Furthermore, supplementation with GlcNAc under glucose-depleted conditions elevated CD109 levels, paralleling the pattern observed for STAT3 activation (Fig. 6D). Moreover, recombinant CD109 is sufficient to activate STAT3 in RKO cells, which have low basal STAT3 activity (Fig. 6E). However, high concentrations of recombinant CD109 were required to stimulate STAT3 activation, suggesting that additional co-factors or cellular context may be necessary to achieve robust STAT3 signaling.

**Figure 6.**
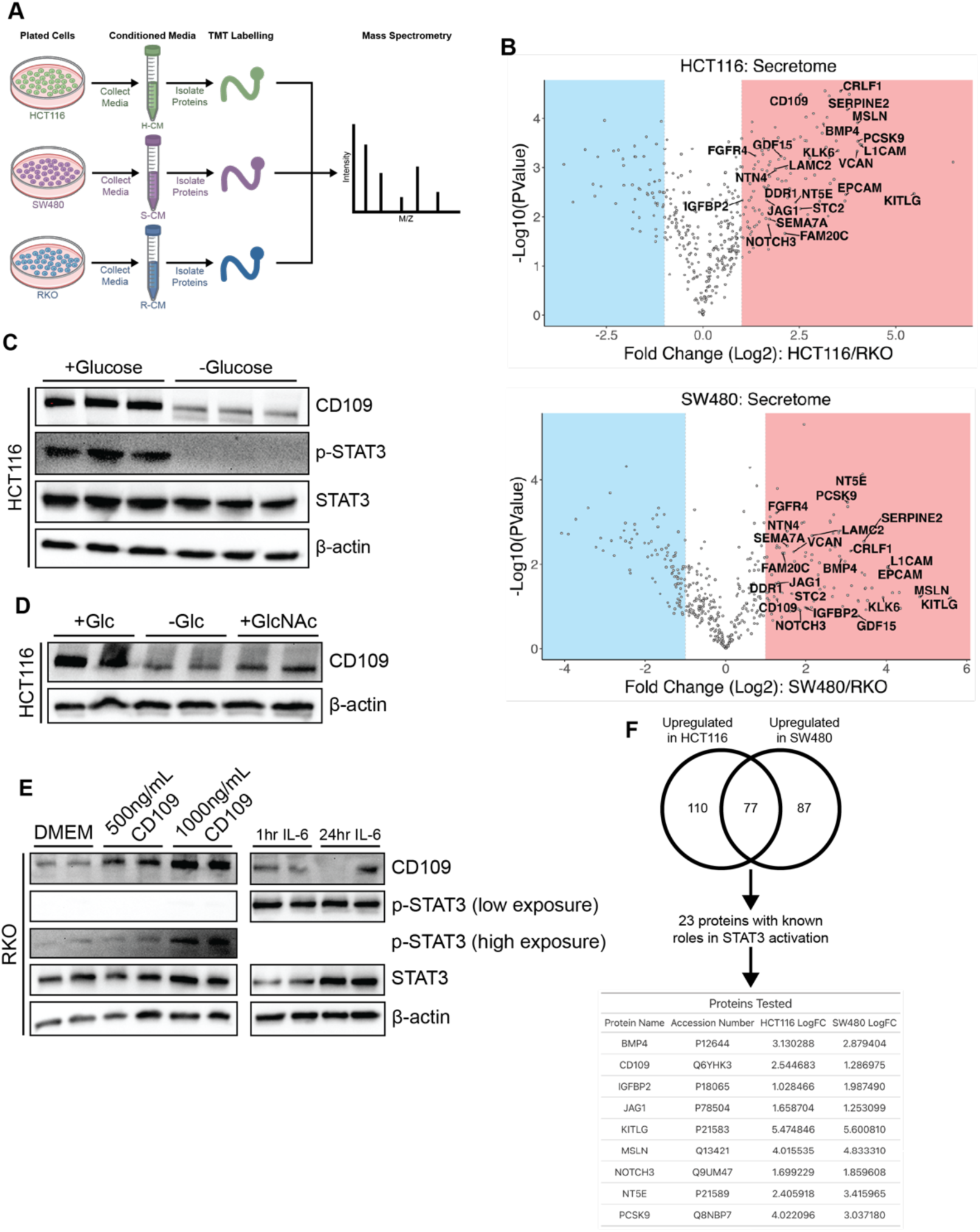
Investigating the identity of the secreted protein. (A) Schematic of the tandem mass tagging (TMT) proteomics experiment. (B) Volcano plots for differentially regulated proteins in the proteomics. The 23 proteins with known roles in STAT3 activation are labelled. (C) HCT116 cells treated with glucose free media for 24 hours to measure levels of CD109. (D) HCT116 cells glucose deprived overnight with 25mM GlcNAc added for 24 hours to measure CD109 levels. (E) RKO cells treated with 500ng/mL or 1000ng/mL CD109 for 24 hours or 20ng/mL IL-6 for 1 hour. (F) Venn diagram of the proteomics data with the 8 proteins tested in the table.

Revisiting the list of 23 proteins, we tested an additional 8 based on fold change and availability of selective inhibitors or active recombinant proteins (Fig. 6F). One protein, KITLG (SCF) was tested by treating glucose deprived HCT116 and SW480 cells with recombinant SCF (Supp. Fig. 6A). Three proteins: BMP4, IGFBP2, and MSLN were tested by treating RKO cells with recombinant protein. No increase in STAT3 activation was observed (Supp. Fig. 6B-D). The remaining four proteins: PCSK9, NOTCH3, JAG1, and NT5E, were tested by inhibition in HCT116 cells, but no decrease in STAT3 activation was detected (Supp. Fig. 6E-G). These findings suggest that while individual proteins may not fully account for the glucose-dependent maintenance of STAT3 activation, they provide important insight into potential candidates, and further investigation may reveal synergistic interactions or the involvement of untested proteins in this process.

### Glucose regulates STAT3 activation in vivo

STAT3 activation is essential for colon cancer growth, however this has been mainly assessed in a colon cancer model that requires intestinal injury and inflammation when cytokine levels are highly induced^8^. To directly assess the requirement for STAT3 in CRC, we generated STAT3 knockout MC38 cells using CRISPR-Cas9 (Supp. Fig. 7A). Tumors derived from STAT3- deficient cells were smaller than those from control cells (Supp. Fig. 7B-C). These findings underscore the critical role of STAT3 activation in CRC progression. To assess the role of STAT3 activation in tumors in vivo, we used MC38 cells, a syngeneic colon cancer cell line that grow in wild-type C57BL6/J mice. MC38 cells exhibit high basal STAT3 activation, like human CRC cell lines (Fig. 7A), and show reduced STAT3 activation upon glucose depletion (Fig. 7B). Consistent with our in vitro findings syngeneic tumors derived from MC38 cells also displayed high STAT3 activation in vivo (Fig. 7C), which is reduced in tumors from mice treated with 2-DG, a glycolytic inhibitor (Fig. 7D). To gain insight into the link between high glucose and STAT3 activation in patients, we used 19 STAT3 target genes weighted equally with GLUT1 gene expression, which revealed lower overall survival when STAT3 target genes and GLUT1 expression are higher (Fig. 7E). Our working model suggests that a group of glycosylated, secreted proteins mediate autocrine and paracrine STAT3 activation in CRC cells (Fig. 7F).

**Figure 7.**
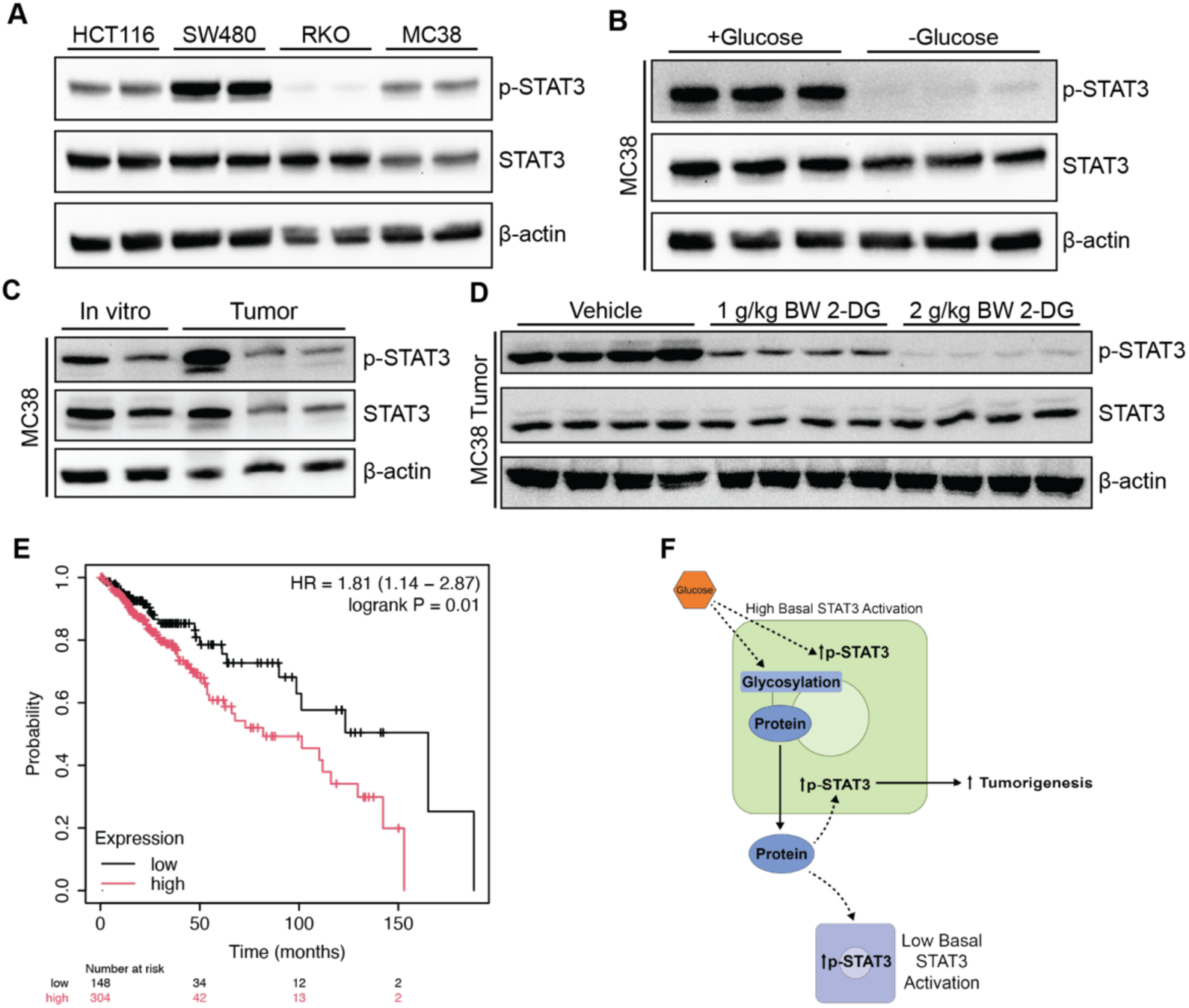
STAT3 activation is high in tumors and regulated by glucose. (A) Western blot of basal pSTAT3 levels in panel of CRC lines, HCT116, SW480, RKO are all human CRC lines. MC38 is a mouse CRC cell line. (B) Western blot of MC38 cells +/- glucose for 24 hours. (C) Western blot measuring basal STAT3 activation in MC38 cells grown in vitro or in vivo tumors. (D) Western blot of activated STAT3 levels from syngeneic tumors for mice treated with vehicle (saline), 1g/kg, or 2g/kg 2-DG for 1 week. (E) Kaplan-Meier plot showing overall survival of patients expressing GLUT1 and STAT3 target genes. (F) Working model of glucose-mediated STAT3 activation.

## DISCUSSION

STAT3 activation is a key driver of CRC progression^8,11–13^. Our data reveal that glucose plays a critical role in sustaining STAT3 activation with the HBP being essential for this process via N-glycosylation, an important post-translational modification commonly found on secreted proteins. Our results suggest that glucose-dependent secreted proteins are essential in maintaining STAT3 activation in CRC and further validate the role of CD109 in STAT3 activation. Additionally, we demonstrate the importance of glucose for STAT3 activation in vivo using a syngeneic mouse model.

STAT3 activation is persistently elevated in CRC^11–13^. However, in canonical STAT3 signaling negative feedback mechanisms, such as SOCS3, typically suppress STAT3 activation^9,10^. IL-6, a cytokine that is commonly elevated in the tumor microenvironment activates STAT3 by binding to its receptor, composed of IL-6 receptor and GP130^7,9^. Previous studies have shown that IL-6 can sustain STAT3 activation through EGFR binding to GP130^40^. However, we observed high levels of STAT3 activation in vitro even in the absence of IL-6. While elevated glucose has been linked to increased STAT3 activation in various cancers, the underlying mechanism remains unclear^41–44^. Our findings support the idea that high glucose levels contribute to sustained STAT3 activation in CRC.

Sustained STAT3 activation has also been associated with pyruvate kinase M2 (PKM2), a glycolytic enzyme that converts phosphoenolpyruvate (PEP) to pyruvate. Knockout and rescue experiments have shown that the PKM2 dimer functions as a protein kinase, whereas the tetramer form acts as an active pyruvate kinase. Both in vitro and cell line studies suggest that PKM2 can directly phosphorylate and activate STAT3 using PEP as a phosphate donor.

Mutants of PKM2 that preferentially form dimers enhance STAT3 signaling and stimulate cell proliferation^45^. However, conflicting evidence exists regarding the role of PKM2 in regulation of STAT3. Hosios et al. challenged the hypothesis that PKM2 functions as a protein kinase, finding no evidence of protein phosphorylation using cancer cell lysates and [^32^P]-PEP^46^. Our research aims to clarify the role of glycolysis in sustaining STAT3 activation. We demonstrate that GlcNAc, a metabolite in the HBP, can activate STAT3 in the absence of glucose, highlighting the importance of glycosylation, and not PKM2, in maintaining STAT3 activation by glucose.

Glycosylation is an essential post-translational modification for many secreted proteins. To identify proteins of interest, we conducted proteomic analysis and identified 77 upregulated proteins in HCT116 and SW480 (high basal activated STAT3) cells compared to RKO (low basal activated STAT3) cells. Of these, 23 have been previously implicated in STAT3 activation. Our findings suggest that CD109 plays a role in maintaining STAT3 activation. However, the activation of STAT3 with CD109 addition was not to the full extent we would expect. We individually tested several other proteins using inhibitors or recombinant proteins. However, none of the individual proteins we tested appeared solely responsible for sustaining STAT3 activation, leaving open several possibilities for further investigation. It is possible that key proteins were not identified in our mass spectrometry analysis, or that multiple proteins act cooperatively to maintain activation. Additionally, many of the identified proteins can bind ligands, lipids, or metabolites that may contribute to STAT3 activation, interactions that would not be detected using our current approaches.

In CRC, STAT3 activation plays an important role in tumor progression^8^. Despite numerous clinical trials, no JAK-STAT3 inhibitor has yet been approved for CRC treatment^6,14^. Our study uncovers a glucose-dependent mechanism for sustaining STAT3 activation, suggesting potential new therapeutic strategies. Diabetes, which is associated with an increased risk of CRC and poor outcomes, may be an important factor in this context^47,48^. Combining therapies to control systemic glucose levels with IL-6-JAK-STAT3 inhibitors could improve patient outcomes. In summary, we have identified a novel glucose-dependent signaling pathway that sustains STAT3 activation in CRC. These findings suggest that other oncogenic pathways may also be influenced by nutrient availability, extending beyond traditional nutrient-sensing mechanisms.

## MATERIALS AND METHODS

### Animals and treatments

Experiments were conducted with 6 to 8-week-old C57BL/6 wild-type male and female mice. The syngeneic model used MC38 empty vector (sgEV) or STAT3 knock out (sgSTAT3) cells. Cells were diluted in DMEM to a concentration of 1x10^7^ cells per mL and 100µL (1 million cells) were injected in each flank. After 3 weeks mice were sacrificed, and tumors were collected and weighed. For in vivo glucose metabolism alteration, mice were injected with MC38 cells, as described above, once tumors reached 500mm^3^ then mice were treated daily with vehicle (saline), 1g/kg, or 2g/kg 2-DG for 1 week. Mice were euthanized three hours after treatment on the final day and tissues were collected for analysis.

### Cell culture and treatments

All cell lines were maintained at 37°C, 5% CO2 in 25mM glucose DMEM supplemented with 10% FBS and 1x antibiotic/antimycotic.

*Nutrient Depletion:* For amino acid deprivation, cells were cultured in amino-acid free DMEM supplemented either with all 13 essential amino acids or with all except glutamine/GlutaMAX and incubated for 24 hours. FBS depletion was performed by culturing cells in DMEM containing amino acids but without FBS for 24 hours. Glucose depletion was carried out either by culturing cells in glucose-free DMEM for 24 hours or overnight, followed by subsequent treatments in a temporal manner.

*Cytokine and Metabolic Treatments:* Cells were treated with 20ng/mL IL-6 (Biolegend, 570802) for 1 hour unless otherwise indicated. Glycolysis was inhibited by treating cells with 2.5 mM 2- DG (Cayman Chemical, D-3179) for 2 hours. Cells were supplemented with 25mM pyruvate overnight or 50mM methyl-pyruvate for 4 hours.

*ER Stress Induction:* ER stress was induced using 6µM tunicamycin (Cayman Chemical, 22038) or 2µM thapsigargin (Cayman Chemical, 10522) for 24 hours.

*Supplementation and Inhibitor studies:* Cells were supplemented with 25mM GlcNAc for 24 hours, unless otherwise stated. Protein trafficking was inhibited using 10ng/mL brefeldin A (Sigma-Aldrich, B5936-200UL) or a 1:1000 dilution of golgi stop (BD, 554724) for 5 hours. *Conditioned Media Experiments:* Cells were treated with a 1:1 mixture of conditioned media and fresh DMEM for 24 hours, unless otherwise specified. The conditioned media was then refreshed using the same dilution for an additional 24 hours before harvesting.

*Recombinant Protein Treatments:* Cells were treated with recombinant CD109 (R&D Systems, 4385-CD-050) at 500ng/mL or 1000ng/mL for 24 hours. For IGFBP2 treatment, 100ng/mL recombinant IGFBP2 (R&D systems, 674-B2; provided by Narla lab) was applied for 1 hour.

Cells were treated with 1µg/mL recombinant mesothelin (MSLN; R&D Systems, 3265-MS-050) for 12 hours. BMP4 treatments were performed with 50µg/mL recombinant BMP4 (PeproTech, 120-05) for 20 minutes. Recombinant Human SCF (KITLG; R&D Systems, BT-SCF) was used at 10 or 100ng/mL for 24 hours in the presence or absence of glucose.

*Small Molecule Inhibitors:* To assess the role of JAK signaling, glucose-deprived cells were treated with 3µM Ruxolitinib (Selleckchem, S1378) for 2 hours before glucose readdition.

Similarly, 100µM Galloflavin (LDH inhibitor; MedChem Express HY-W040118) was used for 2 hours before glucose readdition. Glycosylation was inhibited using 1 or 10µM NGI-1 (MedChem Express, HY-117383) or 50µM OSMI-1 (Cayman Chemical, 21894) for 24 hours. Cells were treated with 5, 15, or 30µM SBC-115076 (PCSK9 inhibitor; MedChem Express HY-12402) for 24 hours. NOTCH signaling was inhibited with 40µM DAPT (EMD4biosciences; provided by Samuelson lab) for 24 hours. Adenosine metabolism was blocked with 10µM AB928 (MedChem Express HY-129393) or 10µm AB680 (MedChem Express HY-125286) for 24 hours.

*Hypoxia Experiments:* Cells were exposed to hypoxia (2% O2) with or without glucose readdition for 4 hours.

*Sugar Supplementation:* To examine the role of different sugars, cells were glucose-deprived overnight and then supplemented with glucose, fructose, galactose, mannose, lactose, or sucrose at concentrations of 5 or 25mM for 24 hours.

### KO cell lines

Knock out cell lines (STAT3 KO MC38) were generated using CRISPR-Cas9 in the LentiCRISPRv1 (Feng Zhang; Addgene plasmid 49535) vector, as previously described^49^. The single guide RNA (sgRNA) sequences used to make KO are listed in Table 1.

### Luciferase

The SIE-luciferase plasmid (pGL4.47 [luc2P/SIE/Hygro]) was purchased from Promega. The SIE-Luc was transfected into HEK293T cells. Additionally, the HEK293T cells were transfected with EV or a JAK1, JAK2, or JAK3 overexpression construct, the JAK overexpression constructs were generous gifts from the Carter-Su lab. The co-transfection was done using polyethylenimine (PEI; Polysciences Inc. 23966), as previously described^50^. Cells were treated with or without glucose for 24 hours prior to being lysed with reporter lysis buffer (Promega).

The luciferase activity was measured and normalized to β-galactosidase.

### Western Blotting

Cells were lysed with RIPA buffer supplemented and samples were run on SDS-polyacrylamide gels and transferred to PVDF membrane. Antibodies from cell signaling technology: phospho- STAT3 Y705 (9145S), STAT3 (9139S), p-AKT S473 (4060S), Akt (4691S), p-P38 T180/Y182 (4511S), P38 (9212S), p-S6 S240/244 (2251S), S6 (2217S), p-SMAD1/5 S463/465 and S465/467 (13820P), pEGFR (3777T), SRC (2108), pERK (9101), pACC (3661), pAMPK (2531), HRP-conjugated anti-rabbit (7074S), HRP-conjugated anti-mouse (7076S). Primary antibodies from Proteintech: β-actin (66009-1-Ig, 1:2000). Primary antibodies from Santa Cruz: GAPDH (sc-47724), p-STAT3, Y705 (sc-136193), STAT3 (sc-293151), ERK (sc-135900), p-SRC (sc-166860), EGFR (sc-373746). All primary antibodies were used at 1:1000 dilution unless otherwise indicated.

### RNA-Seq

RNA-sequencing was performed using the Illumina HiSeq (4000) sequencer with single-end 50- cycle reads. The analysis was performed as previously described^51,52^. The differential expression was tested between no glucose and 4 hours glucose within the cell lines. A cut off false-discovery rate-adjusted (FDR) P value <0.05 was used for marking genes differentially expressed.

### qPCR

RNA was isolated and qPCR was run as previously described using SYBR green dye (Invitrogen 4385616)^53,54^. The primers for qPCR are listed in Table 1. Analysis was done using GraphPad Prism with 2-way Anova.

### Proteomics

*Sample Preparation:* HCT116, SW480, and RKO cell were plated and allowed to adhere overnight prior to changing to serum free media. After 24 hours the conditioned media was collected. Proteins were precipitated by adding sodium deoxycholate and trichloroacetic acid and the pellet was washed 2 times with acetone prior to resuspending in RIPA buffer^55,56^

*TMT-Labelled LC/MS:* Samples were prepared and run as previously described^57^, samples were proteolyzed and labelled with TMT 6-plex (ThermoFisher). Labeled samples were mixed, and dried using a vacufuge. An offline fractionation of the combined sample (∼200 µg) into 8 fractions was performed using high pH reversed-phase peptide fractionation kit according to the manufacturer’s protocol (Pierce; Cat #84868). Fractions were dried and reconstituted in 9µL of 0.1% formic acid/2% acetonitrile in preparation for LC-MS/MS analysis. The replicates of HCT116, SW480, and RKO CM were labeled with TMT channel 126 and 129, 127 and 120, and 128 and 131, respectively. LC/MS analysis was done using multinoch-MS3^58^. Orbitrap Fusion (Thermo Fisher Scientific) and RSLC Ultimate 3000 nano-UPLC (Dionex) was used to acquire the data. Two µL of the sample was resolved on a PepMap RSLC C18 column (75 µm i.d. x 50 cm; Thermo Scientific) at the flow-rate of 300 nl/min using 0.1% formic acid/acetonitrile gradient system (2-22% acetonitrile in 150 min;22-32% acetonitrile in 40 min; 20 min wash at 90% followed by 50 min re-equilibration) and directly spray onto the mass spectrometer using EasySpray source (Thermo Fisher Scientific). Mass spectrometer was set to collect one MS1 scan (Orbitrap; 120K resolution; AGC target 2x10^5^; max IT 100 ms) followed by data-dependent, “Top Speed” (3 seconds) MS2 scans (collision induced dissociation; ion trap; NCE 35; AGC 5x10^3^; max IT 100 ms). For multinotch-MS3, top 10 precursors from each MS2 were fragmented by HCD followed by Orbitrap analysis (NCE 55; 60K resolution; AGC 5x10^4^; max IT 120 ms, 100-500 m/z scan range). Proteome Discoverer (v2.4; Thermo Fisher) was used for data analysis. MS2 spectra were searched against SwissProt Human protein database 20291 entries; reviewed; downloaded on 12/13/2021) using the following search parameters: MS1 and MS2 tolerance were set to 10 ppm and 0.6 Da, respectively; carbamidomethylation of cysteines (57.02146 Da) and TMT labeling of lysine and N-termini of peptides (229.16293 Da) were considered static modifications; oxidation of methionine (15.9949 Da) and deamidation of asparagine and glutamine (0.98401 Da) were considered variable. Identified proteins and peptides were filtered to retain only those that passed ≤1% FDR threshold. Quantitation was performed using high-quality MS3 spectra (Average signal-to-noise ratio of 10 and <50% isolation interference). Secreted proteins were further validated inHCT116 and SW480 conditioned media that was fractionated using a 30kDa molecular weight cut off filter and prepared as above.

*Analysis:* Proteins from the TMT-labelled dataset were filtered to only include proteins in the LC-Tandem MS dataset to ensure the proteins over 30kDa were used for the rest of the assessment as these are implicated in STAT3 activation. The resulting proteins were filtered for those known to be glycosylated (https://glycosmos.org/glycoproteins/) and those that have sequences predicting membrane binding, secretion, or both (https://www.proteinatlas.org/humanproteome/tissue/secretome#what_is_a_secreted_protein).

The remaining proteins were then filtered for a fold change (calculated comparing HCT116 and SW480 to RKO) of greater than or equal to 2. Analysis was done in Python 3 and graphs were generated using R 4.4.1.

### Kaplan-Meier Analysis

KMPlot web-based survival analysis tool tailored for medical research was accessed on 5/9/25^59,60^. The RNA-seq colon cancer dataset was accessed. For analysis, a gene signature for STAT3 activation was accessed to find STAT3 targets that contained the annotated motif of ‘NGNNATTTCCSGGAARTGNNN’ from MSigDB^61,62^. This revealed 22 genes; 19 of these genes were available for querying in the colon cancer RNA-seq dataset including: APBA1, ASXL1, BTBD1, CCL2, CISH, CLDN5, ELMO1, GEN1, HNRNPR, ICAM1, IRF1, MAFF, PROS1, SDHAF2, SERPING1, SLC38A5, TRAF4UBR1, VIP. Analysis was based on mean-expression value where each of these genes was given a weight of 1 and the gene for GLUT1 (SLC2A1) was given an equal weight to the totality of the 19 STAT3 genes such that the end analysis equally weight GLUT1 expression and STAT3 activation. The other following parameters were used in this analysis - Survival: Overall survival, Auto select best cutoff: percentile, Follow up threshold: all, Start follow-up at: all, Compute median over entire database: FALSE, Cutoff value used in analysis: 3768.85, Expression range of the probe: 1692 - 48213, Probe set option: user selected probe set, Proportional hazards assumption: checked, analysis not restricted by Pathology, gender, race, or treatment status. Generated P value: 0.0103; hazard ratio 1.81.

## Supporting information

Supplemental References

## Authorship Contributions

KB, KT, XX, and YMS conceived and designed the study; KT, SKR, and XX developed the methodologies; KB, KT, CM, NA, SM, HNB, JANC, HG, VP, SKR, and XX acquired the data; KB, KT, SK, HNB, SKR, CAL, XX, and YMS analyzed and interpreted the data; KB, KT, XX, and YMS supervised the study and wrote the manuscript; and all authors edited and provided input to the manuscript.

---

**Supplemental Figure 1.**
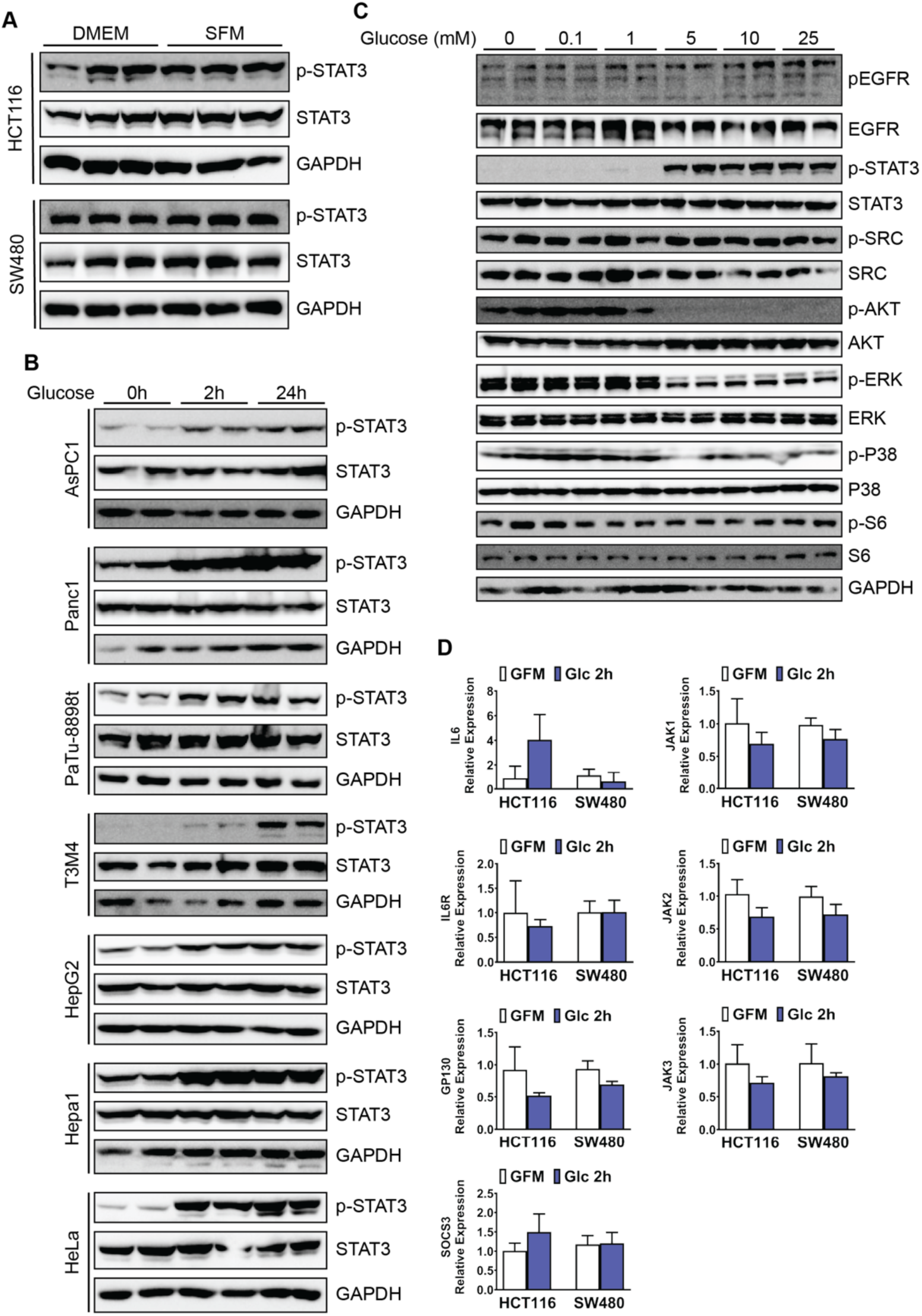
S**T**AT3 **activation is regulated by glucose in many cancers.** (A) HCT116 and SW480 cell lines treated with DMEM supplemented with 10% FBS or DMEM with no FBS for 24 hours. (B) Human pancreatic cancer cell lines (AsPC1, Panc1, PaTu-8898t, T3M4), human hepatocellular carcinoma cell line (HepG2), murine hepatocellular carcinoma cell line (Hepa1), and human cervical cancer cell line (HeLa) treated with glucose (25mM) for 0, 2, or 24 hours. (C) HCT116 cells treated with increasing glucose concentrations for 24 hours. (D) HCT116 or SW480 treated with 25mM glucose for 0 or 2 hours prior to harvesting RNA to measure gene expression by qPCR.

**Supplemental Figure 2.**
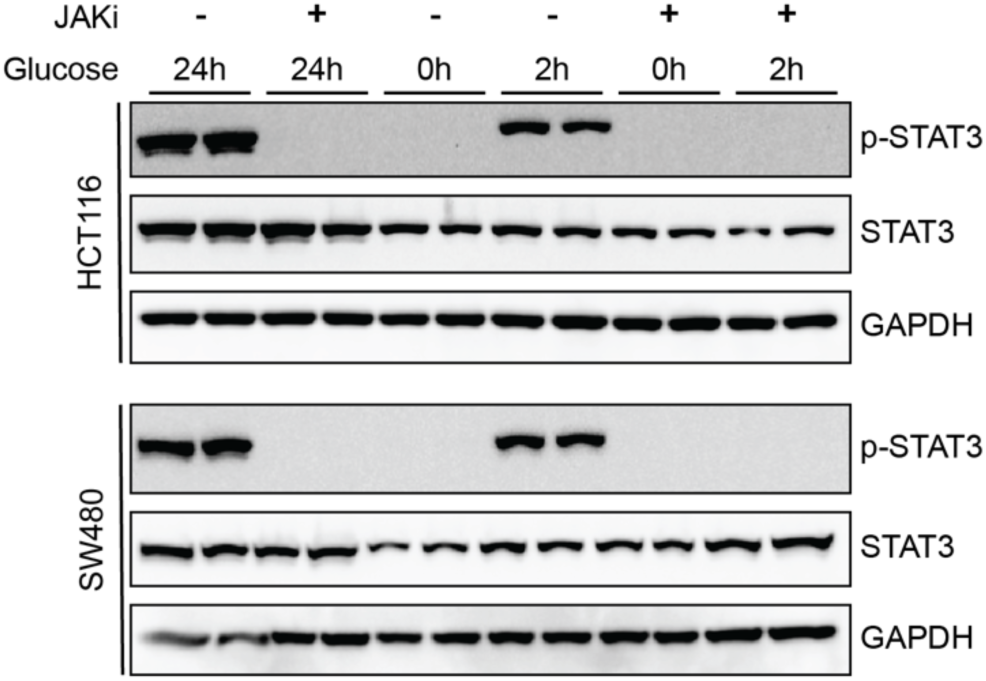
G**l**ucose**-dependent STAT3 activation is JAK dependent.** HCT116 and SW480 cell lines treated with Ruxolitinib (3µM) for 2 hours then glucose (25mM) for 0, 2, or 24 hours.

**Supplemental Figure 3.**
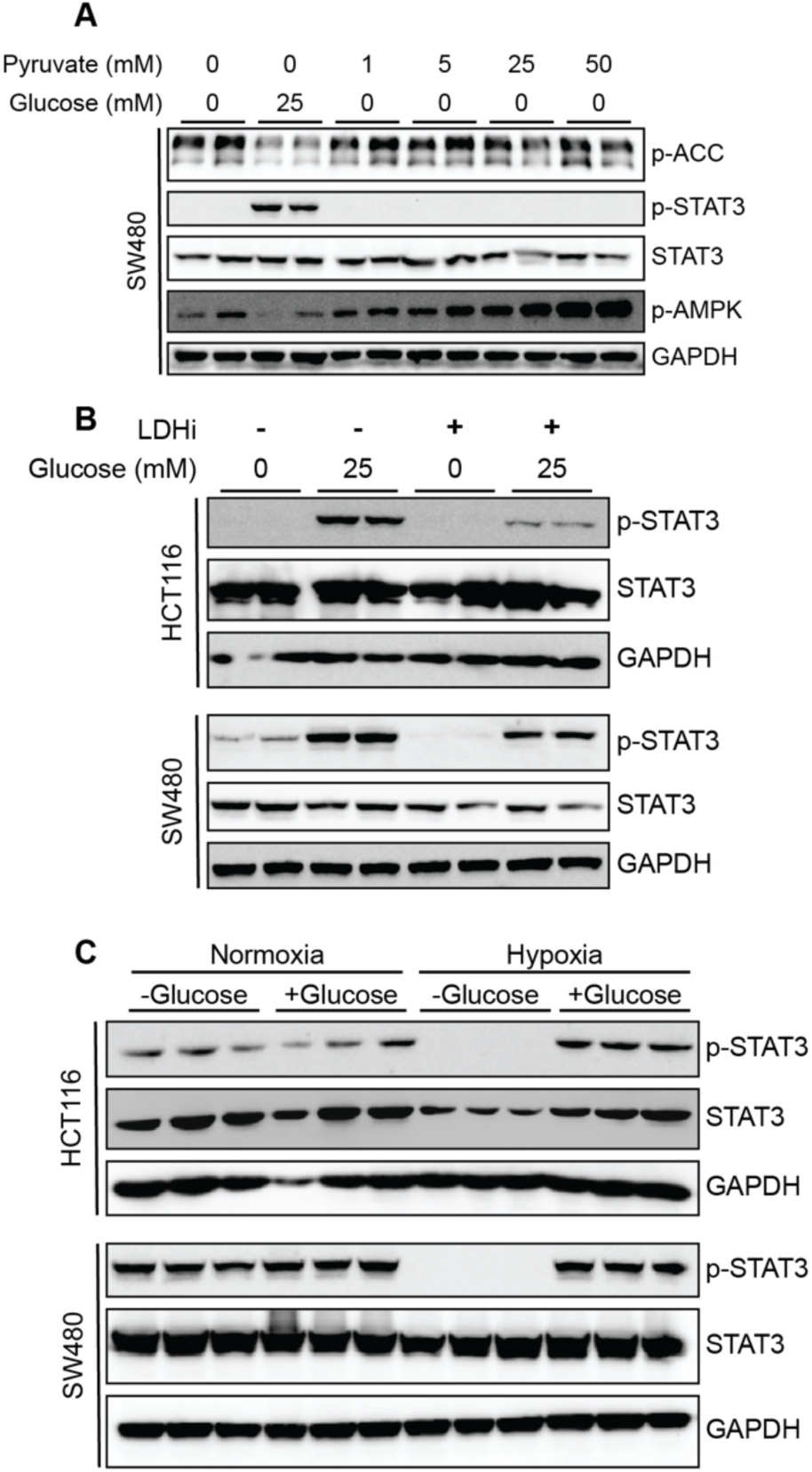
G**l**ucose **metabolism and hypoxia effects on STAT3 activation.** (A) SW480 cell line treated with 0 or 25mM glucose and increasing concentrations of sodium pyruvate overnight. (B) HCT116 and SW480 cell lines treated with 100µM Galloflavin (LDH inhibitor) for 2 hours then glucose (25mM) for 0, 2, or 24 hours. (C) HCT116 and SW480 cells treated with or without glucose and placed in hypoxic conditions (2% oxygen) for 4 hours.

**Supplemental Figure 4.**
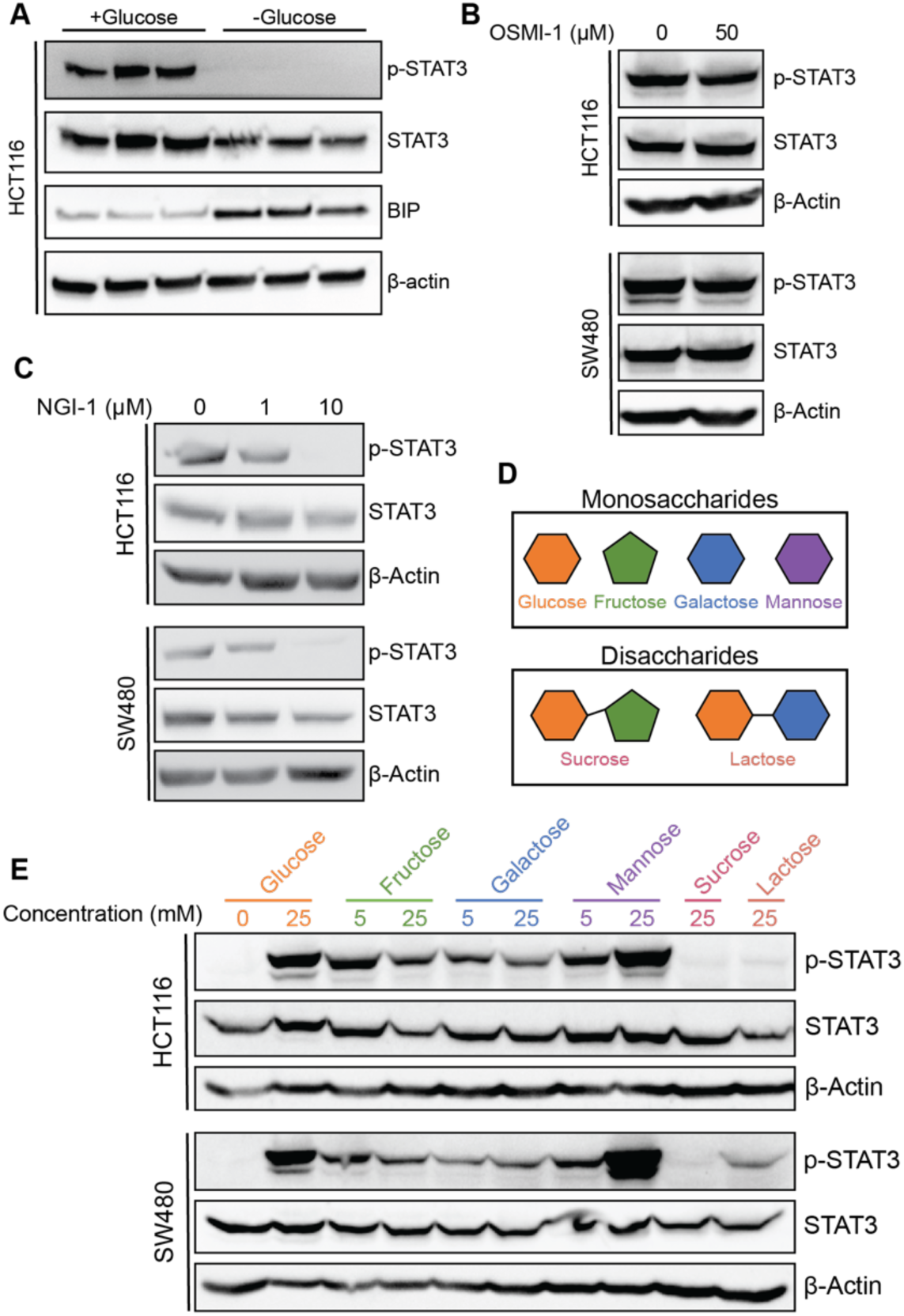
Glycosylation and sugar metabolism effects on STAT3 activation. (A) HCT116 cells treated with or without glucose (25mM) for 24 hours to measure BIP levels as an indicator of ER stress. (B) HCT116 and SW480 cells treated with 0 or 50 µM OSMI-1, O-GlcNAc transferase inhibitor, for 24 hours. (C) HCT116 and SW480 cells treated with 0, 1, or 10µM NGI-1, an oligosaccaryltransferase inhibitor, for 24 hours. (D) Schematic of sugars used to treat cells. (E) HCT116 and SW480 cell lines treated with 0 or 25mM glucose and glucose free media with 5 or 25mM fructose, galactose, mannose, or 25mM sucrose or lactose for 24 hours.

**Supplemental Figure 5.**
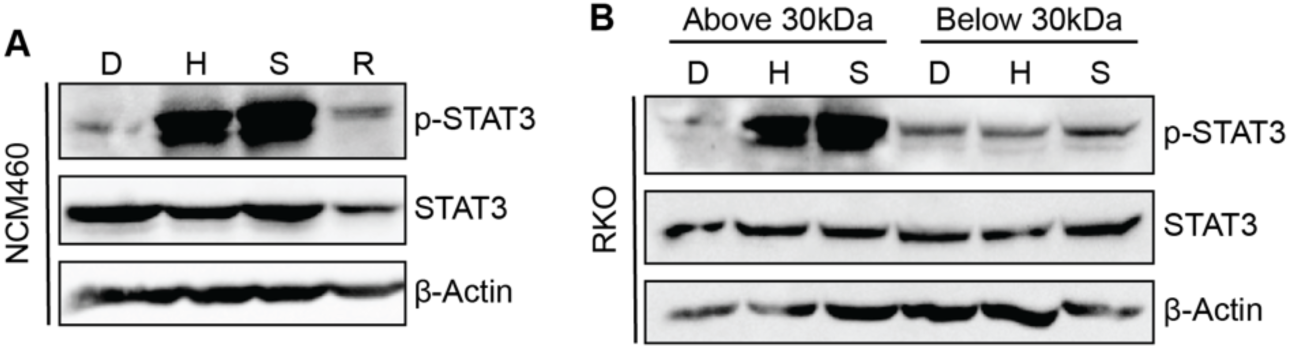
H**C**T116 **and SW480 conditioned media activate STAT3 in other cells lines.** (A) NCM460 cell line (normal colon epithelium cell line) treated with DMEM {D}, HCT116 condition media {H}, SW480 condition media {S}, or RKO condition media {R} for 48 hours with a refresh of the condition media at 24 hours. (B) RKO cells treated with DMEM {D}, HCT116 condition media {H}, or SW480 condition media {S} that was passed through a 30kDa molecular weight cut off filter for 48 hours with a refresh of the condition media at 24 hours.

**Supplemental Figure 6.**
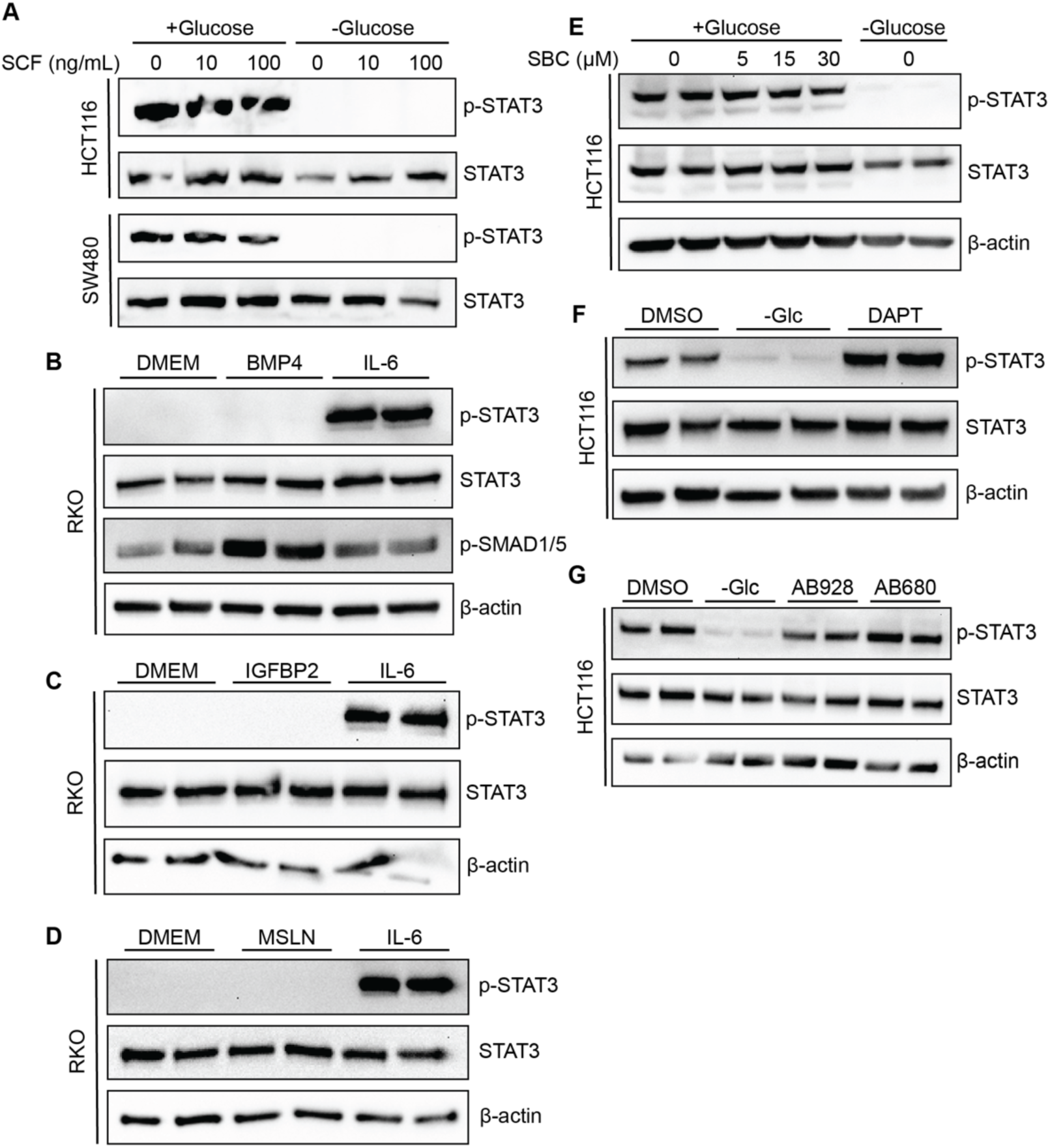
I**n**hibition **of proteins or treatment with recombinant proteins identified in proteomics.** (A) Western blot showing HCT116 and SW480 cell lines treated with varying concentrations of recombinant SCF (KITLG) for 24 hours in the presence or absence of glucose. (B) Western blot showing RKO cells treated with 50ug/mL BMP4 recombinant protein for 20 minutes or 20ng/mL IL-6 for 1 hour. (C) Western blot showing RKO cells treated with 100ng/mL IGFBP2 recombinant protein for 1 hour or 20ng/mL IL-6 for 1 hour. (D) Western blot showing RKO cells treated with 1µg/mL MSLN recombinant protein for 12 hours or 20ng/mL IL- 6 for 1 hour. (E) Western blot showing HCT116 cells treated with varying concentrations of SBC-115076 (SBC), a PCSK9 inhibitor for 24 hours. (F) Western blot showing HCT116 cells treated with 40µM DAPT, a Notch inhibitor for 24 hours. (G) Western blot showing HCT116 cells treated with 10µM AB928, an adenosine receptor inhibitor, or 10µM AB680, a NT5E inhibitor for 24 hours.

**Supplemental Figure 7.**
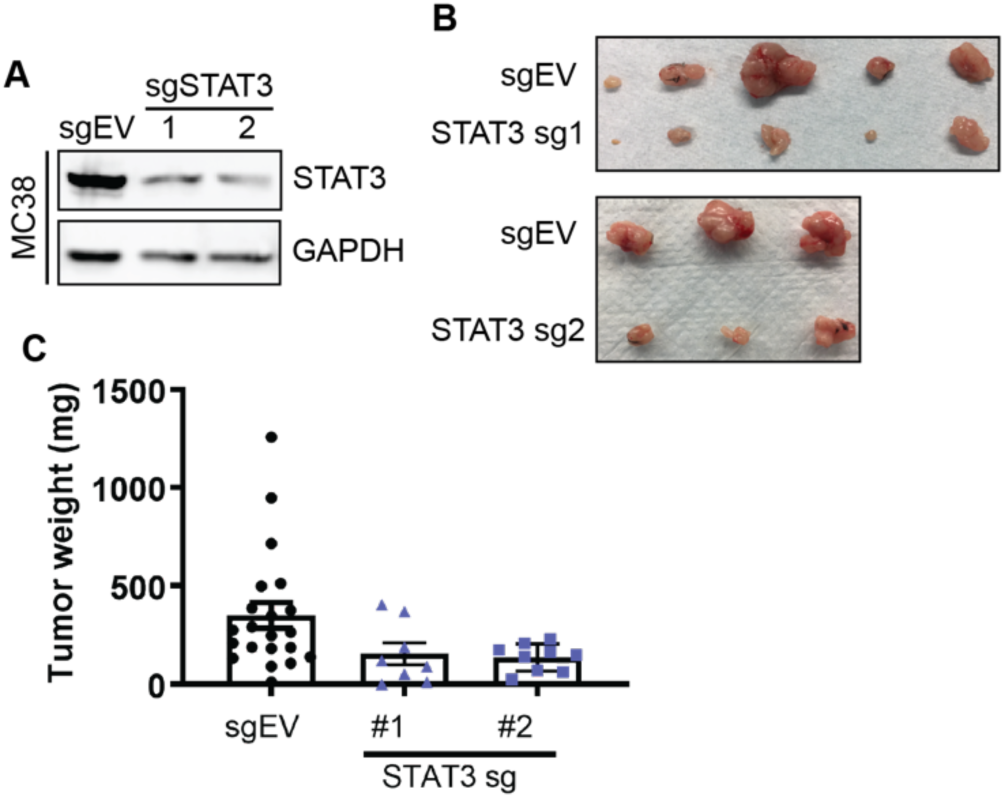
S**y**ngeneic **tumor model with STAT3 knockout MC38 cells.** (A) Western blot of STAT3 levels and GAPDH loading control for MC38 cells transfected with empty vector (EV) or STAT3 single guide RNA (sg) to generate CRISPR-Cas9 knockouts of STAT3. (B) Tumors from MC38-EV or MC38-STAT3 knockout syngeneic mouse model. (C) Tumor weight of MC38-EV or MC38-STAT3 knockout syngeneic mouse model. P value was determined using one-way Anova.

## Supplemental Materials

**Supplemental Table 1.** RNA-Seq data with fold changes between no glucose and 4 hours glucose exposure.

**Supplemental Table 2.** Proteomics data of 77 proteins upregulated in HCT116 and SW480 cells compared to RKO cells, annotated with known roles in STAT3 activation (with citations)

**Supplemental References.** Used in Supplemental Table 2 for protein roles in STAT3 activation.

